# Integrative Analysis of Gene Expression, Protein Abundance, and Metabolomic Profiling Elucidates Complex Relationships in Chronic Hyperglycemia-Induced Changes in Human Aortic Smooth Muscle Cells

**DOI:** 10.1101/2024.08.28.610153

**Authors:** Smriti Bohara, Atefeh Bagheri, Elif G. Ertugral, Igor Radzikh, Yana Sandlers, Peng Jiang, Chandrasekhar R. Kothapalli

## Abstract

Type 2 diabetes mellitus (T2DM) is a major public health concern with significant cardiovascular complications (CVD). Despite extensive epidemiological data, the molecular mechanisms relating hyperglycemia to CVD remain incompletely understood. We here investigated the impact of chronic hyperglycemia on human aortic smooth muscle cells (HASMCs) cultured under varying glucose conditions *in vitro*, mimicking normal (5 mmol/L), pre-diabetic (10 mmol/L), and diabetic (20 mmol/L) conditions, respectively. Patient-derived T2DM-SMCs served as disease phenotype controls. Results showed significant increases in cellular proliferation, area, perimeter, and F-actin expression with increasing glucose concentration (*p* < 0.01), albeit not exceeding the levels in T2DM cells. Atomic force microscopy analysis revealed significant decreases in Young’s moduli, membrane tether forces, membrane tension, and surface adhesion in SMCs at higher glucose levels (*p* < 0.001), with T2DM-SMCs being the lowest among all the cases (*p* < 0.001). T2DM-SMCs exhibited elevated levels of pro-inflammatory markers such as IL-6, IL-8, and MCP-1 compared to glucose-treated SMCs (*p* < 0.01). Conversely, growth factors such as FGF-2 and TGF-β were higher in SMCs exposed to 10 mmol/L glucose but lower in T2DM-SMCs (*p* < 0.01). Pathway enrichment analysis showed significant increases in the expression of inflammatory cytokine-associated pathways, especially involving IL-10, IL-4 and IL-13 signaling in genes that are up-regulated by elevated glucose levels. Differentially regulated gene (DGE) analysis showed that compared to normal glucose receiving SMCs, 513 genes were upregulated and 590 genes were downregulated in T2DM-SMCs; fewer genes were differentially expressed in SMCs receiving higher glucose levels. Finally, the altered levels in genes involved in ECM organization, elastic fiber synthesis and formation, laminin interactions, and ECM proteoglycans were identified, which highlight the role of hyperglycemia in vascular remodeling and CVD progression. Our results collectively suggest that chronic hyperglycemia in vascular SMCs leads to morphological, biomechanical, and functional alterations, potentially contributing to the pathogenesis of T2DM-associated CVD. The observed differences in gene expression patterns between *in vitro* hyperglycemic models and patient-derived T2DM-SMCs highlight the complexity of T2DM pathophysiology and underline the need for further studies.

## Introduction

T2DM is a serious public health concern and a global burden. It is one of the leading causes of mortality affecting more than half a billion people worldwide, and directly contributing to at least a few million deaths annually^1^. T2DM detection is difficult in its early stages leaving many people undiagnosed, and often independently leads to the onset of CVD by the time of diagnosis^2–4^ T2DM is a chronic metabolic and inflammatory condition, marked by inadequate insulin secretion by pancreatic β-cells and gradual buildup of tissue insulin resistance, with significant implications in accelerated vascular aging and CVD^5, 6^. T2DM is a complex multifactorial disease and has been implicated in the onset of coronary and peripheral artery diseases (e.g., atherosclerosis, stroke), microvascular (e.g., retinopathy, nephropathy) and neuropathic complications, hypertension, dyslipidemia, and arterial stiffness^7^.

When the insulin secretion in diabetic patients cannot maintain glucose homeostasis, it results in hyperglycemia, hyperinsulinemia, inflammation, and higher body fat content, which in turn dysregulate the immune system, adipokines, gut microbiota, and a host of other tissue functions, resulting in CVD^8^. Despite its prevalence and the availability of epidemiological data related to T2DM pathophysiology, the molecular mechanisms by which hyperglycemia contributes to CVD are still being elucidated^9, 10^. In healthy vasculature, ECs play a crucial role in maintaining vessel homeostasis and VSMC contractile phenotype by releasing numerous factors (e.g., VEGF, NO). T2DM conditions lead to endothelial dysfunction, release of various inflammatory markers (e.g., ROS, interleukins, TNF-α), and elevated oxidative stress that disturbs this homeostasis and activates SMCs, resulting in vasoconstriction, thrombus formation, and the onset of CVD^11–14^. When the functioning EC layer is compromised, VSMCs are exposed to high glucose levels in their microenvironment and the glucose molecules could easily pass through their cell membrane leading to intracellular hyperglycemia and their dedifferentiation^15–17^. Elucidating the role of chronic higher glucose levels on VSMC phenotype and genotype could offer insights into the fundamental mechanisms by which hyperglycemia progresses and offer therapeutic avenues for modulating undesirable outcomes.

The direct contribution of any singular factor on T2DM complications is hard to isolate in both humans and animal models, especially that of hyperglycemic levels. VSMCs cultured *in vitro* in high glucose conditions (≥ 20 mmol/L) exhibited proliferation, migration, adhesion, necrosis via H_2_O_2_ formation, and elevated secretion of various biomolecules (e.g., ROS, TNF-α, MMP-2), along with activation of numerous pathways (e.g., JNK, ERK, AKT, NF-κB, MAPK)^18–23^. In these cultures, cells were typically exposed to high glucose levels for a short duration (3 h – 72 h), or rodent SMCs were used in lieu of human cells, which do not efficiently mimic the chronic buildup of high blood glucose levels in humans *in vivo*. The potential benefits of numerous drugs tested in such studies to regulate VSMC behavior *in vitro* or in animal models *in vivo* have not led to breakthroughs in clinical trials, suggesting that a deeper understanding of the co-expression between metabolomic, transcriptomic, and proteomic transformations in these cells under prolonged exposure to elevated glucose concentrations mimicking pre-diabetic and diabetic conditions is critical to delineate the specific role of high glucose levels in T2DM initiated CVD.

In this study, we investigated the role of hyperglycemic conditions on the activation of VSMC pathways that contribute to early onset of endothelial dysfunction and CVD. We cultured human aortic smooth muscle cells (HASMCs; isolated from multiple donors) for 21 days to broadly mimic normal (5 mmol/L glucose), prediabetic (10 mmol/L glucose), and diabetic (20 mmol/L glucose) states. The phenotypic and biomechanical changes in cells under these conditions were assessed using AFM and confocal imaging. Metabolomic, proteomic, and RNA-seq analysis were performed to compare with those from human T2DM−HASMCs. Our study reveals the biological pathways activated during early stages of T2DM and identifies crucial biomarkers triggered during hyperglycemic conditions.

## Methodology

### Cell cultures

Cryopreserved HASMCs from multiple patients (lot 70008916, isolated from 44-year old Caucasian female; lot 2164581, isolated from 44-year old Caucasian female; lot 1917076, isolated from 40-year old Caucasian female) and T2DM−HASMCs (lot 3125, isolated from tunica media of fibrous plaques-free aorta of 49-year old type 2 diabetic Caucasian female) were purchased from American Type Culture Collection (ATCC, VA, USA), Thermo Fisher Scientific (Waltham, MA, USA), and CELL Application Inc. (San Diego, CA, USA). The cell culture medium was prepared using normal D-glucose (5 mmol/L)^24^ added to Dulbecco’s Modified Eagle Medium (L-DMEM) supplemented with 10% heat-inactivated fetal bovine serum (FBS), and 1% penicillin/ streptomycin. All cell culture medium supplies were from Thermo Fisher Scientific.

For all the experiments, primary HASMCs and T2DM−HASMCs cells were initially passaged in T-25 culture flasks (Nunc^TM^, Thermo Fisher Scientific) coated with 0.1% gelatin. Prior to experiments, cells were serum starved in L-DMEM (containing 1% heat-inactivated FBS, 1% pen-strep) for 24 h to synchronize cell cycles. Then, all HASMCs were exposed to varying D-glucose levels (5 mmol/L, 10 mmol/L, 20 mmol/L) conditions^25^, and T2DM−HASMCs were exposed to 5 mmol/L D-glucose levels for 21 days in 6−well plates coated with 0.1% gelatin (n = 6 wells/ condition/ assay). To mimic diabetic conditions *in vitro*, VSMCs were initially cultured in normal glucose (5 mmol/L) levels for at least 72 h, before switching to high glucose (> 20 mmol/L) levels for the rest of the study^26^. The glucose concentration in the medium was measured using the GlucCell^TM^ glucose monitoring system (CESCO Bioengineering Co., Trevose, PA, USA). The final cultures were harvested and processed for biomechanical testing using AFM, transcriptomic, proteomic, and metabolic data analysis omics analysis, and cytokines/ chemokines/ growth factors quantification.

### Phenotype assessment

Separately, SMCs (5 × 10^4^ cells/well) were seeded into 24-well plates and cultured under various glucose conditions for up to 7 days. At the end of each time point (days 1, 4, 7), 100 µL of MTT reagent (Sigma Aldrich) was added to each well and incubated for 1 h (37 °C, 5% CO_2_). Then, 100 µL of detergent added to each well, incubated for 1 h at room temperature, and the absorbance measured at 570 nm with a BioTek Synergy H1 Hybrid multi-mode microplate reader. Wells with complete media alone and no cells were processed in similar way. The fold-change in cell density was calculated by normalizing to the readout from the initial seeding density. SMCs under various glucose conditions as well as T2DM cells were imaged at random locations using a Zeiss Axiovert A1 fluorescence microscope equipped with Hamamatsu camera and image acquisition software. At least ten images per condition were analyzed using NIH ImageJ software to quantify the perimeter and area of the cells.

### Metabolomic analysis

After 21 days of culture, the medium was removed while maintaining the plates on ice, and the cell layers were sequentially washed with cold 1× PBS and cold deionized (DI) water. The cell layers were then incubated for three minutes with methanol containing 5% acetic acid at 4 °C and scraped using a cell scraper. The cell layers were pooled in a 15 mL centrifuge vial, placed on a cell rocker for 30 min in a cold room, and sonicated on ice for 15 sec at 0.5 pulse rate (Branson Digital Sonifier; Marshall Scientific, Hampton, NH, USA). Metabolites were extracted by vortexing the samples for 30 sec at high speed, followed by centrifugation at 17,000 ×*g* for 10 min at 4 °C. The supernatants were carefully transferred to autosampler vials and analyzed immediately by HPLC-MS untargeted conditions.

LC/QTOF analysis was performed with Agilent 6545 QTOF Mass Spectrometer coupled with Agilent 1290 Infinity II UHPLC system (Agilent Corp., Santa Clara, CA, USA). Chromatographic separation was achieved with Waters XSelect HSS T3 (2.1 mm × 100 mm, 2.1 µm) column. The mobile phase was composed of water/0.1% formic acid (A) and acetonitrile/0.1% formic acid (B). The gradient elution conditions for both positive and negative modes are as follows: 0-6 min, 5% B; 6-8 min, 35% B; 8-30 min, 90% B; 30-35 min, 5% B; 35-37 min, 5% B. Post-run equilibration time was set to 5 min, flow rate set to 0.3 mL/min, and column temperature set to 35 °C. Untargeted polar metabolomics data was acquired with Agilent MassHunter Data Acquisition software (Version B.10.1.48). Source parameters were set as follows: drying gas (N_2_) temperature 300 °C and flow 11 L/min, sheath gas (N_2_) temperature 350 °C and flow 11 L/min; nebulizer gas (N_2_) pressure 40 psi; capillary voltage 2000 V; nozzle voltage 0 V; fragmentor voltage, 70 V; skimmer voltage, 30 V; The data obtained was processed using molecular feature extraction and time alignment using Agilent MassHunter Profinder 10.0. A 100% in at least one sample group feature presence was used to remove irreproducible features. Extracted molecular features were imported into Agilent Mass Profiler Professional (MPP) and Metaboanalyst for identification and statistical analysis. Statistical thresholds of *p* < 0.05 and fold of change (FC) > 1.5 were chosen for selecting significant molecular features. Identification of molecular features was performed utilizing both METLIN database and in-house built libraries with mass deviation < 5 ppm. Identified compounds validation was performed using internal standards and recursive analysis of pooled QC sample.

### Proteomics analysis

After 21 days of culture (n = 6 wells/ condition), cell layers were harvested on an ice platform and centrifuged at 1.5 *rcf* for five minutes at 4 °C. Cell count was done using a mini automated cell counter (ORFLO Technologies, Ketchum, ID, USA). The cell supernatant solution was discarded, and the cell pellets were processed to extract the intracellular and extracellular proteins using the Compartmental Protein Extraction Kit (Millipore Sigma) protocols recommended by the vendor. The volume for each buffer used in our study was calculated based on cell count (3.8 ×10^6^ to 4 ×10^6^ cells). We separated cytoplasmic (C), nuclear (N), membrane (M), cytoskeleton (CS), and extracellular matrix protein (ECM) from each well (2 independent repeats). After the extractions, the samples were homogenized, and the protein concentrations measured were between 7.17 µg to 15 µg. The samples thus obtained comprise buffers that were not suitable for direct digestion and LC-MS/MS analysis. Hence, the samples were fractionated on an SDS-page gel, each gel lane was cut into five bands, the bands were digested with trypsin, and the digests were analyzed by LC-MS/MS (Thermo Scientific Orbitrap Exploris™ 480 Vanquish Neo UHPLC). LC-MS data was searched against the human SwissProtKB database using the program Sequest. Spectral count and peptide molecular weight was used to estimate the relative abundance of proteins in these samples. More than 5400 proteins were identified in the C samples (e.g., GAPDH, actin, myosin-9, pyruvate kinase, tubulin, filamin-A, filamin-C), and over 4900 proteins were identified in the CS samples (e.g., vimentin, myosin-9, actin, pyruvate kinase PKM, filamin-C, GAPDH). The final normalized data were calculated in log_10_-fold change to determine the magnitude of change in protein abundance between the experimental groups. Such changes were used to identify up- and down-regulation of genes. Furthermore, pathway enrichment analysis for differentially expressed genes using ReactomePA^27^ enabled identification of biological pathways that are differentially regulated.

### Cytokine/ chemokine/ growth factors analysis

The cytokines, chemokines and growth factors released in HASMC and T2DM−SMCs cultures were quantified using Discovery Assays® (Eve Technologies, Alberta, Canada) as we detailed earlier^28^. Briefly, the pooled spent media from each culture condition at day 21 were processed using Human Cytokine/ Chemokine 71-Plex Discovery Assay® Array (sensitivity: 0.5 – 10 pg/mL) and Human MMP & TIMP Panel assay (sensitivity: 0.5–14 pg/mL). Analytes were quantified using multiplex LASER bead technology and bead analyzer (Bio-Plex 200), where antibody-conjugated fluorophore beads simultaneously detect multiple analytes from a single assay.

### RNA isolation and sequencing

RNA was extracted from cultured HASMCs and T2DM−HASMCs using Trizol RNA isolation reagent protocol (Thermo Fisher Scientific). Briefly, cell layers were washed with cold 1× PBS, and 1 mL of Trizol reagent was added to each well, and cells were harvested using a cell scraper. Cells were homogenized using a sonicator (Branson Digital Sonifier) and incubated at room temperature for five minutes. A 200 µL aliquot of chloroform was added to the cell supernatant and the colorless aqueous phase containing RNA was separated carefully without disturbing interphase. To ensure that each library meets the required standards for sequencing, the library quantity and quality are evaluated using Tapestation and Qubit. Tapestation (Agilent 4200 Tapestation System) high-throughput quality control device was used for RNA fragment analysis and a 2100 Agilent Bioanalyzer was used for checking RNA integrity number (RIN) score (that ranged from 9.6 − 9.8). Further, a Qubit Fluorometer (Invitrogen) was used to measure accurate concentration of RNA samples. All the samples were pooled, and molarity for sequencing is determined via qPCR (QuantaBio Q qPCR). Finally, the pooled libraries were sequenced using Novaseq 6000 platform with Illumina’s sequencing reagents. Data processing and quality control checks were performed on raw fastq files. Subsequently, the readings were aligned with bowtie2 and quantified using RSEM (RNA-Seq by Expectation-Maximization). Differential expression analysis was conducted on mapped read counts obtained from RSEM using the R/Bioconductor package EBSeq. The analysis identified upregulated and downregulated genes, which were then used for Gene Set Enrichment Analysis and Venn diagram plotting.

### Confocal microscopy

HASMCs under three different conditions mimicking healthy, prediabetic and diabetic conditions, and T2DM−HASMCs were cultured for 21 days in multi-chamber cell culture slides (Thermo Fisher Scientific). The cell culture medium was discarded, and cells were washed with 1× PBS, fixed with 4% paraformaldehyde for 30 min, permeabilized with 0.1% Triton X-100 in PBS for 5 min, and blocked with 2% bovine serum albumin (BSA) for 30 minutes. The cells were washed with 1× PBS and incubated with phalloidin-iFluro 488 (abacam) and 4′,6-diamidino-2-phenylindole (DAPI, Sigma), at room temperature for 90 min, to stain for F-actin and nuclei, respectively. Cells were washed with 1× PBS to remove excess staining solution and imaged using a Nikon A1Rsi confocal laser scanning microscopy. To quantify actin expression in SMCs exposed to varying glucose levels, images were captured with a 40× objective, transformed into grayscale images, and mean fluorescence intensity of actin filaments analyzed using Fiji ImageJ software. Three independent experiments were performed. Several fields of view of equal size per glucose treatment were randomly selected and mean fluorescence intensity of actin expression were measured and compared to controls.

### Atomic force microscopy

HASMCs were cultured for 7 days in 60−mm gelation-coated petri dishes under various glucose concentrations, while T2DM−HASMCs were cultured under normal glucose conditions. Cells were maintained at 37 °C for live-cell nanoindentation performed using an MFP-3D-Bio AFM (Asylum Research, Oxford Instruments, Santa Barbara, CA) mounted on an inverted optical microscope (Nikon Eclipse Ti). Tip-less AFM cantilevers (Nano World, Arrow TL 1, nominal spring constant ∼ 0.03 N/m) were modified by attaching a 5-μm polystyrene bead using epoxy. The actual spring constant was determined using the thermal calibration method in a clean culture dish before each experiment and thermal/mechanical equilibrium of AFM was achieved by submerging probe in cell medium for stabilization. For each condition, at least 50 cells were randomly chosen and intended at random locations at 0.3 Hz with constant trigger point of 0.3 V. Force-indentation curves were obtained and Young’s modulus (*E_Y_*) was determined from these curves using Hertz’s contact model. The force of adhesion (*F_ad_*) and tether forces (*F_T_*) were obtained from the force-indentation curves, from which membrane tension 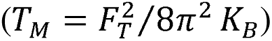 and tether radius 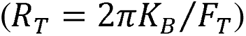 were calculated^29^. Frequency distribution and one-way ANOVA were performed for statistical analysis and *p* < 0.05 was considered significant.

### RNA-seq data analysis

RNA-seq reads were mapped to the human reference genome (version: GRCh38) and corresponding annotated protein-coding genes using Bowtie^30^, allowing up to 2-mismatches. The gene expected read counts and Transcripts Per Million (TPM) were estimated by RSEM^31^. The TPMs were further normalized by EBSeq^32^ R package to correct the potential batch effect. The EBSeq package^32^ was further used to assess the probability of gene expression (mRNAs) being differentially expressed between any two given conditions. We required that DEGs should have FDR < 5% via EBSeq and >2 fold-change of “median-by-ratio normalized read counts +1”.

### Pathway enrichment analysis

The Reactome pathway enrichment analysis was performed by the “ReactomePA” R package^33^.

### Statistical Analysis

Data from human SMCs and T2DM SMCs were pooled and expressed as scatter plots with mean ± SEM where appropriate. Descriptive statistical analyses were conducted using the GraphPad Prism 10 software. A one-way ANOVA with post hoc Tukey’s test was used for multiple comparisons of the results from various assays, with statistical significance set at *p* < 0.05.

## Results and Discussion

### Hyperglycemic conditions induce phenotypic changes in human aortic SMCs

Significant differences in morphology and phenotype were evident in SMCs exposed to various glucose conditions (**Fig. 1, A, B**). It was noted that cells were more aggregated, irregular in shape, and larger in cross-sectional area within high glucose cultures, contrasting with the regular and standalone appearance in 5 mmol/L cultures. T2DM cells lacked the hill and valley configuration seen in normal SMCs, in line with that reported by others^34^. Analysis of the cell morphologies from multiple such images revealed that the cell area and perimeter significantly increased at higher glucose concentration (**Fig. 1, C**; *p* < 0.05), whereas T2DM cells were the largest in size (*p* < 0.05) among all the cases. This is to be expected because high glucose levels reportedly have differential effects on the phenotype and function of many cell types. The cell morphology was analyzed using the cell shape index, i.e., *CSI* = *4πA/P*^2^, where *A* and *P* are the measured cell area and perimeter, respectively.^35^ Results indicated that SMCs exposed to 10 mmol/L and 20 mmol/L glucose had *CSI* around 0.23 ± 0.01, whereas T2DM cells and normal SMCs (5 mmol/L) had CSI around 0.48 ± 0.03 (*p* < 0.001). It should be noted that the area and perimeter of the SMCs in control (5 mmol/L glucose) cultures is in broad agreement with literature^36^. Quantitative analysis revealed elevated levels of F-actin expression in SMCs receiving higher glucose concentrations (**Fig. 1, D, E**). The area covered by actin within the cells as well as the fluorescence intensity of actin were significantly higher in 20 mmol/L glucose treated cells and T2DM-SMCs compared to cells receiving 5 mmol/L or 10 mmol/L glucose (*p* < 0.05). Collectively, these results suggest that exposure of VSMCs to elevated glucose levels for longer periods induces significant phenotypic changes in these cells, broadly comparable to their T2DM counterparts.

**Figure 1.**
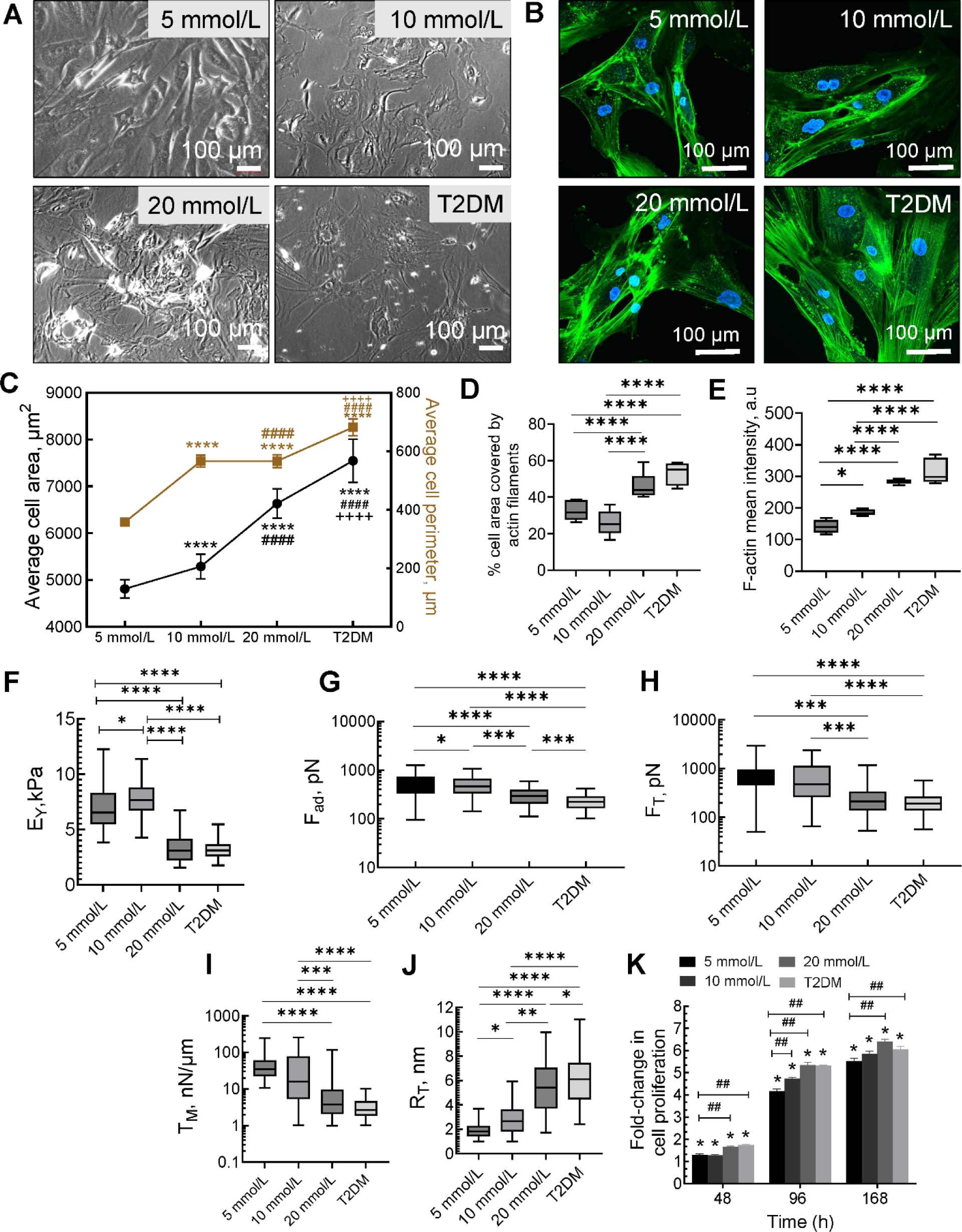
**A**. Representative phase-contrast images of adult human aortic SMCs cultured under varying glucose concentration (5, 10, or 20 mmol/L) and T2DM SMCs. **B**. Representative confocal images of filamentous-actin (F-actin) and nuclei (blue) stained with Alexa Fluro 488-phalloidin and DAPI, respectively. **C**. Significant differences in average cell area and perimeter were noted between SMCs treated with various dosages of glucose and with T2DM SMCs. Data shown represents mean ± standard error in respective cases (n > 50 cells/condition). **** indicates *p* < 0.0001 vs. 5 mmol/L cultures; #### indicates *p* < 0.0001 vs. 10 mmol/L cultures; ++++ indicates *p* < 0.0001 vs. 20 mmol/L cultures. **D**. Quantitative analysis of area occupied by F-actin within SMCs exposed to various culture conditions. **E**. Quantitative analysis of fluorescence intensity of F-actin. Analysis of fluorescence intensity was done at the original magnification by measuring the mean gray value with Fiji ImageJ software. Biomechanical characteristics such as elastic modulus (*E_Y_*, **F**), forces of adhesion (*F_ad_*, **G**), membrane tether forces (*F_T_*, **H**), membrane tension (*T_M_*, **I**), and tether radius (*R_T_*, **J**) were quantified from the AFM data. *E_Y_* data were calculated by applying Hertz model to force–indentation curves (n ≥ 100 cells/condition) obtained from cells. *F_ad_* and *F_T_* were measured by retraction of beaded-AFM probe from the cell surface. In plots D-J, the center line in the box plots denotes the median, and bound of box shows 25^th^ to 75^th^ percentiles, while upper and lower bounds of whiskers represent the maximum and minimum values, respectively. * indicates *p* < 0.05, ** indicates *p* < 0.01, *** indicates *p* < 0.001, and **** indicates *p* < 0.0001. **K**. Proliferation of human SCMs under various glucose conditions as well as that of T2DM-SMCs. * indicates *p* < 0.05 vs. initial cell seeding density, ## indicates *p* < 0.01 for control (5 mmol/L) vs. glucose treatment (10 mmol/L, 20 mmol/L; T2DM).

The Young’s modulus (*E_Y_*) of SMCs in normal glucose conditions (5 mmol/L) was noted as 7.03 ± 2.01 kPa (**Fig. 1, F**). While 10 mmol/L glucose did not contribute to significant changes in *E_Y_* (7.78 ± 1.65 kPa (*p* > 0.05 vs. 5 mmol/L), 20 mmol/L glucose significantly reduced *E_Y_* to 3.32 ± 1.3 kPa (*p* < 0.05 vs. 5 mmol/L; *p* < 0.05 vs. 10 mmol/L glucose). T2DM cells exhibited *E_Y_* (3.19 ± 0.84 kPa) similar to SMCs that received 20 mmol/L (*p* < 0.01 vs. 5 mmol/L; *p* < 0.01 vs. 20 mmol/L). The *E_Y_* of SMCs in 5 mmol/L glucose conditions was similar to our previous study^37^, whereas the *E_Y_* of T2DM-SMCs is in close range to values reported in literature^38^. Human coronary SMCs derived from healthy patients reportedly had higher *E_Y_* than those isolated from T2DM patients^38^, in agreement with our results here. However, in coronary vascular SMC cultures, isolated from control or diabetic mice, no significant impact of glucose concentration on *E_Y_* was reported^38^. SMCs receiving 5 mmol/L glucose recorded adhesion forces (*F_ad_*) of 558 ± 229 pN (**Fig. 1, G**), and increasing the glucose concentration significantly decreased these adhesion forces (*p* < 0.05 vs. 5 mmol/L; *p* < 0.05 for 10 mmol/L vs. 20 mmol/L). T2DM cells had the lowest adhesion forces of all the cases tested (233 ± 71 pN; *p* < 0.05 vs. all other cases). The tether forces (*F_T_*) recorded in SMCs cultured with 5 mmol/L glucose was 787 ± 342 pN (**Fig. 1, H**), and while 10 mmol/L glucose had no effect on these values, 20 mmol/L significantly reduced the *F_T_*(286 ± 109 pN; *p* < 0.05 vs. 5 mmol/L and 10 mmol/L). T2DM cells recorded the lowest *F_T_* among all the test cases and was not significantly different from 20 mmol/L condition.

The force needed to deform a cell membrane, also termed as the apparent membrane tension (*T_M_*), was calculated from tether forces using 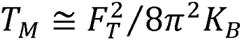, where *K_B_* is the bending stiffness that lies in the 0.1 – 0.3 pN.µm range^39, 40^. The *T_M_* values were as follows (**Fig. 1, I**): 5 mmol/L (50.4 ± 2.3 nN/µm), 10 mmol/L (53.7 ± 3.2 nN/µm), 20 mmol/L (11.4 ± 0.9 nN/µm), and T2DM (3.5 ± 0.2 nN/µm). The *T_M_* was significantly lower in cells receiving 20 mmol/L glucose versus other glucose concentrations (*p* < 0.01), and T2DM cells have the lowest *T_M_* of all the cases tested (*p* < 0.01). The tether radius (*R_T_*) increased with increasing glucose concentration (**Fig. I, J**), with control (5 mmol/L) cells having 1.9 ± 0.03 nm on average and T2DM cells having 6.1 ± 0.13 nm. To the best of our knowledge, the *F_T_*, *F_ad_*, *T_M_* and *R_T_* values for human SMCs and specifically under diabetic conditions haven’t been reported earlier in literature.

Previously, we reported that primary SMCs derived from human aneurysmal aortae and analyzed using AFM exhibited the following values^41^: *E_Y_* = 20.9 ± 7.7 kPa, *F_ad_* = 1.87 ± 0.13 nN, *F_T_* = 218.8 ± 14.3 pN, *T_M_* = 6.07 ± 0.8 nN/µm, and *R_T_* = 2.91 ± 0.19 nm. Compared to T2DM SMCs and SMCs receiving higher glucose in the current study, it is evident that human aneurysmal aortic SMCs (yet another CVD condition) have significantly higher *E_Y_*, *F_ad_* and *R_T_* (*p* < 0.01 in all the cases), although the tether forces and membrane tension of T2DM SMCs were comparable to that in aneurysmal SMCs, underlying the differences in the pathologies of these two vascular diseases.

Compared to SMCs cultured for 24 h, cell proliferation increased 1.3 – 1.75 −fold after 48 h, 4.1 – 5.35 −fold after 96 h, and 5.5 – 6.1 −fold after 7 days (**Fig. 1, K**) in all the cases. Cell proliferation increased with time at each glucose concentration, as well as in T2DM cultures. Our findings are consistent with previous reports where high glucose led to significant increase in the proliferation of various cell types compared to normal glucose levels^42^. For instance, previous studies reported a 3−fold increase in VSMC proliferation exposed to high glucose (> 20 mmol/L) compared to controls (5 mmol/L), as measured using the MTT assay^43–45^.

### Differential effects of hyperglycemia on cytokine/ chemokine/ growth factor release by SMCs

The release of cytokines and chemokines in chronic SMC cultures were quantified (**Fig. 2**) and significant variations in analyte concentrations depending on glucose concentration were noted. Some interleukins that weren’t expressed at all or released in extremely low levels by T2DM cells (e.g., IL-9, IL-17A, IL-17F, IL-28A, IL-4, IL-15, IL-25, IL-33) were seen in SMC cultures receiving glucose (**Fig. 2, A**). On the other hand, interleukins such as IL-6, IL-8, IL-23 and IL-27 were released in higher levels in T2DM cells than in SMCs receiving glucose. Among various cytokines and chemokines tested (**Fig. 2, B**), higher levels of SCD40L, RANTES, GROα, MCP-1, M-CSF, and MIP-1δ were noted in T2DM cultures than in SMC cultures receiving glucose. On the other hand, G-CSF was higher in SMC cultures receiving glucose than in T2DM cultures. There weren’t any significant differences in the expression of other cytokines and chemokines between the various culture conditions. T2DM has been linked to immune system disorders as well as elevated levels of IL-6, IL-18, MCP-1, and TNF-α^46–48^. Although TNF-α, IL-18, and IFN-γ levels were similar in all the cases we tested, IL-6 and MCP-1 levels were significantly higher in T2DM cells consistent with literature.

**Figure 2.**
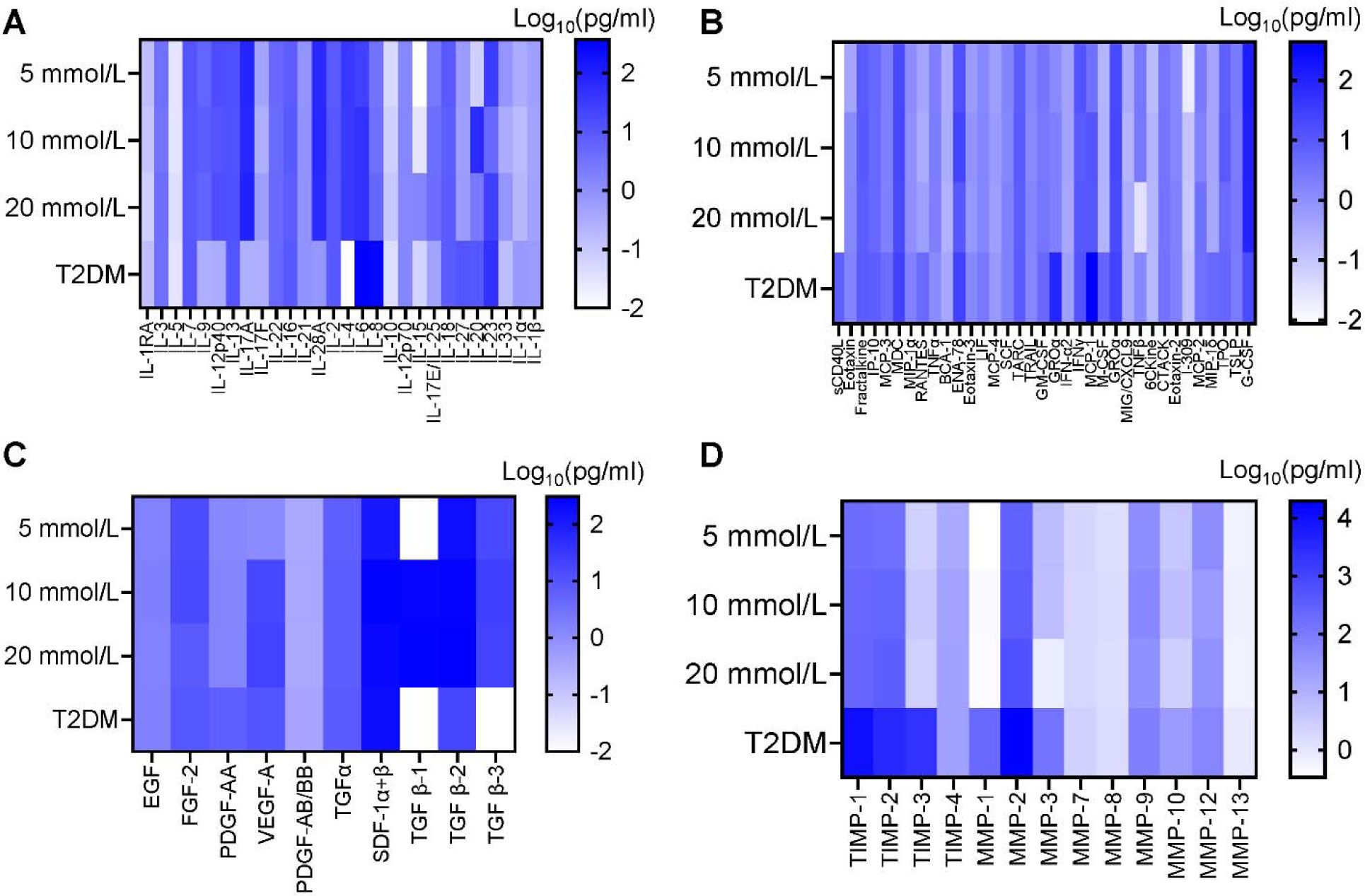
Heat maps of the levels of interleukins (**A**), cytokines & chemokines (**B**), growth factors (**C**), and MMPs/TIMPs (**D**) released in SMC cultures receiving various glucose concentrations and from T2DM cells over 21 days. MMPs–1, 2, 3 and TIMPs–1, 2, 3 were elevated in T2DM cells than glucose-receiving chronic SMC cultures (Fig. 2**, D**). These results suggest that SMCs treated with varying glucose concentrations directly influence the expression of various cytokines, chemokines, growth factors and MMPs/ TIMPs. High glucose exposure was shown to increase transcription and translation of MMPs−1, 2, 9 and 13 in human SMC cultures and their enzymatic activity, especially in the presence of macrophages, representative of T2DM conditions *in vivo*^51^. Consistent with this report, we here note that these specific MMPs were significantly higher in T2DM cultures, whereas MMPs−2, 9 and 12 were also present in SMC cultures receiving high glucose levels. The interplay between MMPs and TIMPs regulate the development of atherosclerotic plaques and vascular ECM remodeling under diabetic conditions^52^. In humans with T2DM and hypertension, elevated levels of TIMPs-1 and 3, TIMP-1:MMP-2 ratio, and TIMP-1:MMP-9 ratio were reported^53, 54^, which mirrors our observations in this study.

FGF-2 levels were slightly higher in SMC cultures receiving 5 and 10 mmol/L glucose, whereas TGF-β2 and TGF-β3 levels were significantly lower in T2DM cultures (**Fig. 2, C**). The levels of VEGF-A, PDGF-AA, and TGF-β1 were higher in cultures receiving 10 and 20 mmol/L glucose. Similar to our observation, high glucose (20 mmol/L) was shown to elevate TGF-β1 and TGF-β-R1 receptor expression in vascular SMCs, via protein kinase C (PKC-α) activation^49^. Hyperglycemia was shown to induce VEGF-A expression in SMCs similar to acute insulin treatment^50^, with implications in decreased function and failure of multiple organs (e.g., retina, kidneys). Our study showed that VEGF-A levels were significantly elevated (> 40-fold; *p* < 0.001 vs. 5 mmol/L) in the presence of 10 mmol/L and 20 mmol/L glucose to levels noted in T2DM cells.

It is worth noting that previous studies suggest that when human aortic SMCs were co-cultured with macrophages under normal (5.5 mmol/L) and high glucose (25 mmol/L) conditions, SMCs exhibited augmented gene and protein expression of MMP-1 and MMP-9, significant increase in MMP-9 enzymatic activity, higher levels of soluble and functionally-active MCP-1 linked to MMPs upregulation, and activated PKCα signaling pathway that together with NF-kB is responsible for MMPs-1, 9 upregulation^51^.

### Inflammatory cytokines-associated pathways are enriched in hyperglycemia induced upregulated genes

Previous research has independently analyzed the transcriptome, proteomics, and metabolomics to identify distinct markers in diabetic SMCs^55^. Our current study, however, profiles these as paired omics data. We explored which genes correspond with metabolomics profiling patterns and the implications of these correlations within the omics modules. We also examined how mRNA levels align with protein abundance. This integrated dataset analysis deepens our understanding of chronic hyperglycemia-induced changes in human aortic smooth muscle cells, advancing beyond mere biomarkers to explore their complex relationships.

We first investigated the genes upregulated by hyperglycemia enriched pathways. Vascular SMCs receiving 5 mmol/L glucose were compared to those receiving 10 mmol/L (**Fig. 3, A**) or 20 mmol/L (**Fig. 3, B**) glucose, and T2DM-SMCs (**Fig. 3, C**). Pathway enrichment analysis was further performed by the differentially expressed genes (DEGs). We found that inflammatory cytokine-associated pathways, especially those involving IL-10 signaling, are prominently enriched in genes that are up-regulated by increased glucose levels of 10 mmol/L and 20 mmol/L glucose, compared to normal glucose levels. IL-10 signaling pathway has been shown to modulate the function of various immune cells. Studies have shown that elevated IL-10 pathways are associated with multiple autoimmune diseases^35^. Additionally, we observed that genes activated by a higher glucose concentration of 20 mmol/L are more closely associated with inflammatory cytokine pathways, including not only IL−10 but also ILs − 1, 4 and 13 signaling pathways (**Fig. 3, B**). This aligns with the hypothesis that long-term exposure to high glucose levels leads to enhanced inflammatory responses, which are associated with chorionic inflammation. The genes involved in the activation of G-alpha signaling events were significantly enriched in SMCs receiving 20 mmol/L glucose but not in SMCs supplied with 10 mmol/L glucose. G-alpha is involved in the inhibition of cAMP dependent pathway which in turn leads to reduced activity of cAMP-dependent protein kinases, as well as activation of the protein tyrosine kinase c-Src^56, 57^.

**Figure 3.**
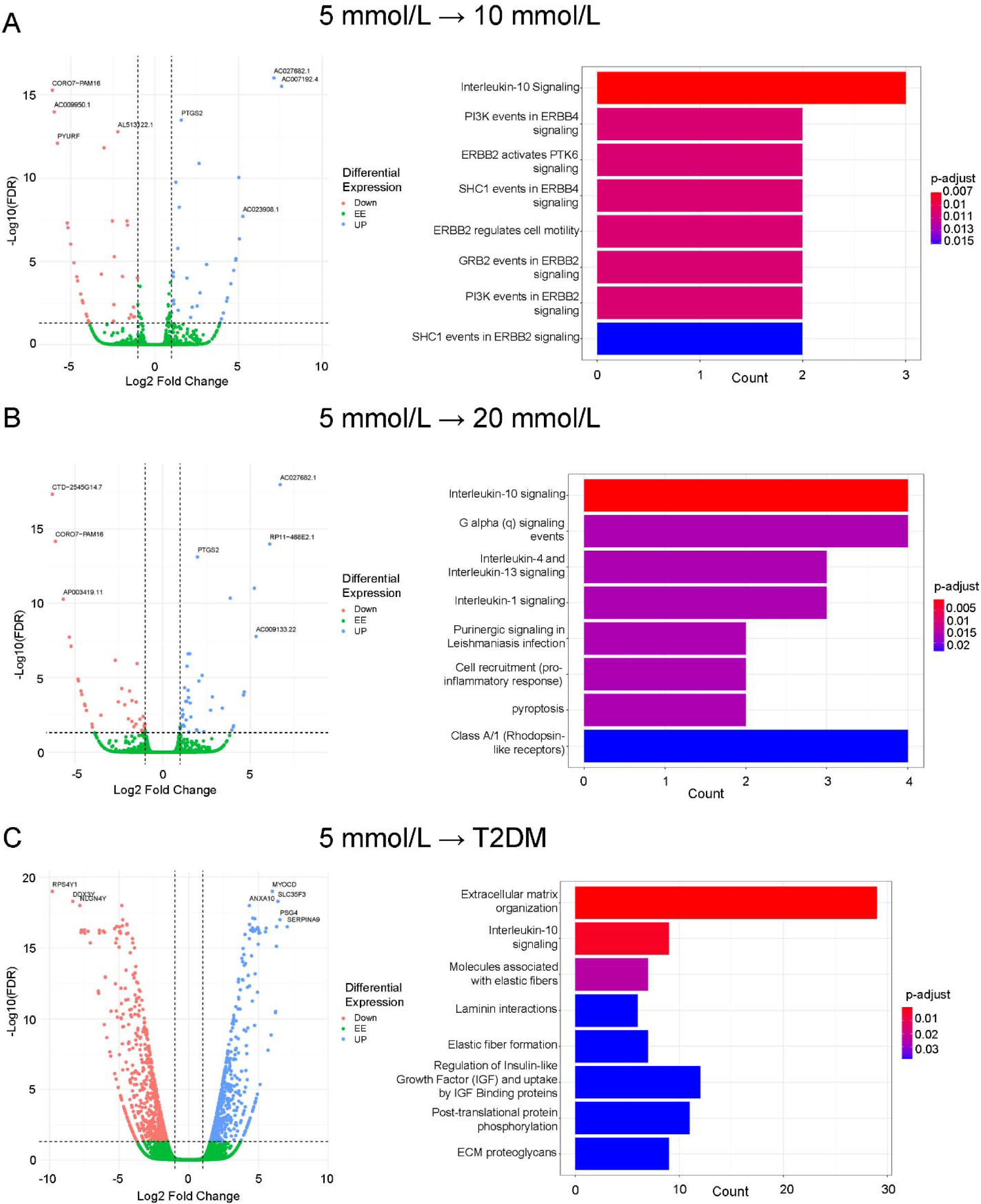
Volcano plots of differential gene expression patterns and enriched pathways in response to glucose concentration and diabetic conditions. (**A**) 10 mmol/L vs. 5 mmol/L glucose, (**B**) 20 mmol/L vs. 5 mmol/L glucose, and (**C**) T2DM Vs 5 mmol/L glucose.

The epidermal growth factor receptor family (EGFR or ErbB 1−4) has been implicated in various cellular functions (e.g., growth, division, differentiation, migration, apoptosis) and multiple downstream signaling pathways (e.g., ERK1/2, MAP, PI3-kinase/Akt) under hyperglycemia or diabetes (types I and II) conditions and related cardiovascular outcomes^58^. In our study, compared to normal glucose levels, chronic exposure to 10 mmol/L but not 20 mmol/L glucose appears to have resulted in differentially expressed genes related to ErbB2 and ErbB4 and their functions and pathways (cell motility, PI3K, PTK6, SHC1) in human vascular SMCs.

We found a notable distinction between the gene expression patterns in cells exposed to high glucose levels *in vitro* (10 or 20 mmol/L) and those compared to T2DM-SMCs (**Fig. 3, C**). Despite some similarities, the up-regulated genes in T2DM cells did not entirely correspond with those observed in the high glucose *in vitro* models. This could possibly be due to the multifactorial nature of T2DM *in vivo* and exposure of SMCs to chronically (multi-year) elevated levels of high glucose and inflammatory molecules under such conditions. In T2DM-SMCs, the dominant pathways involved relate to ECM organization (e.g., laminin, elastin, collagens, proteoglycans, GAGs) and the regulation of IGF, highlighting the complexity of diabetes pathogenesis beyond the changes induced by hyperglycemia alone. This is to be partially expected because T2DM diagnosis could also be an indicator of the onset of various proteolytic vascular conditions such as atherosclerosis that results in vascular ECM remodeling and SMC activation, which is reflected in the cells isolated from a T2DM tissue. Interestingly, IL-10 pathways were enriched in both the *in vitro* models and in T2DM-SMCs, suggesting that increased inflammatory cytokine-associated pathways are a consistent feature of prolonged hyperglycemia. This could potentially contribute to the pathogenesis of diabetic complications, particularly in vascular tissues, implying that controlling inflammation may be crucial in preventing these complications. Interestingly, compared to SMCs receiving normal glucose levels, T2DM cells had altered gene expression for molecules involved in ECM organization, elastic fiber synthesis and formation, laminin interactions, and ECM proteoglycans, highlighting the role of T2DM in vascular remodeling and CVD progression.

Similar outcomes were noted from the differentially regulated gene (DGE) analysis (**Fig. 4**) performed on these cells. We performed statistical analysis on the normalized read count data to assess quantitative changes in expression levels between the various groups. Compared to SMCs receiving normal glucose levels, 513 genes were upregulated, and 590 genes were downregulated in T2DM cells (list of genes provided in **Table S1**). We note that, among others, the genes involved in vascular matrix remodeling were significantly upregulated in T2DM cells (e.g., COL, ELN, GLB, FBLN, LMN). We performed a ReactomePA analysis to identify the pathways enriched for the upregulated and downregulated genes, respectively). Cells receiving 10 and 20 mmol/L glucose have much fewer genes that were differentially expressed, while very few of these genes were common between all the cell types.

**Figure 4.**
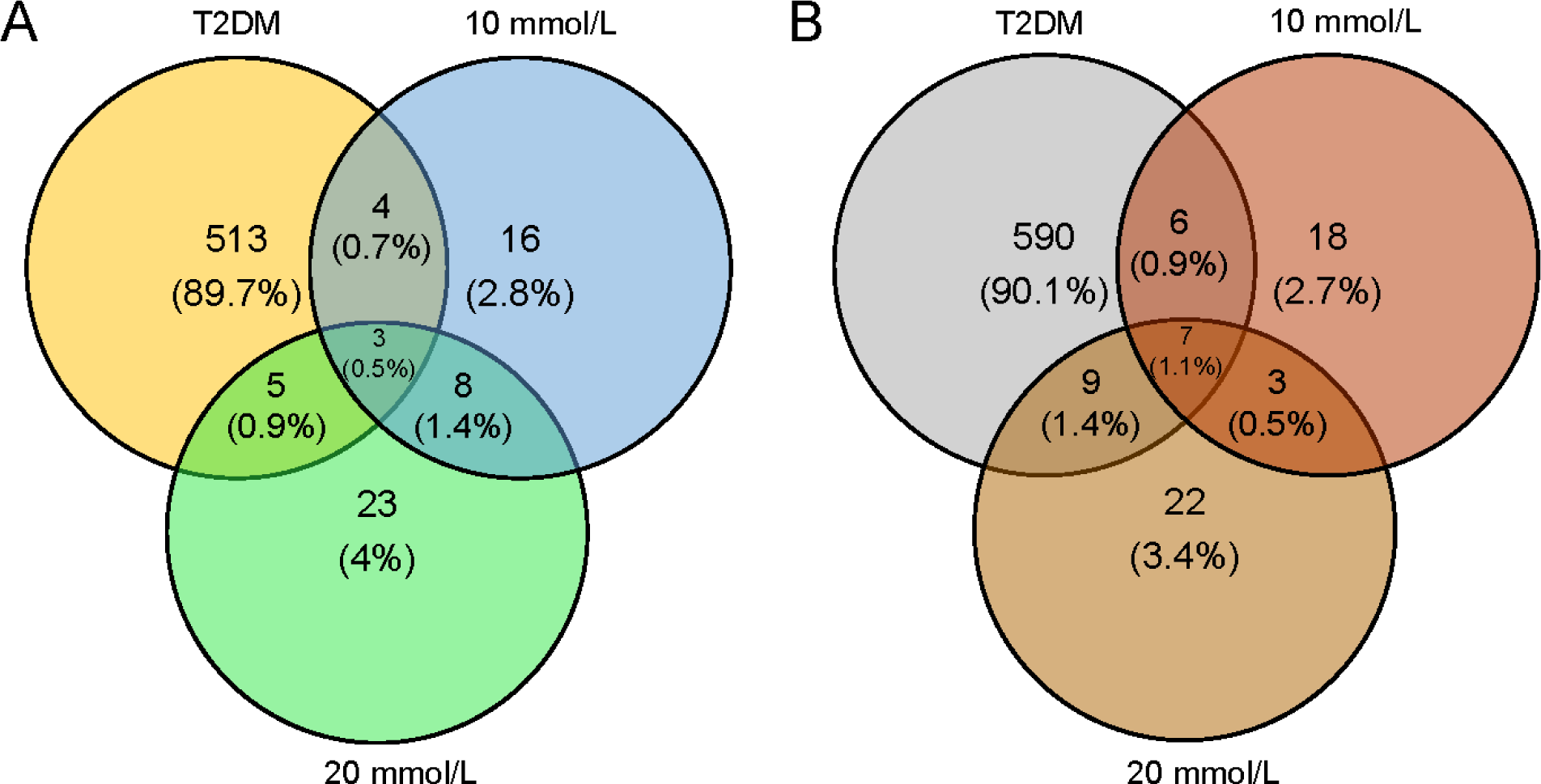
Venn diagram indicating the overlap between differentially expression genes (DEG) in SMCs receiving various glucose concentrations (10 mmol/L and 20 mmol/L). RNA-seq data for each condition was compared to cells receiving 5 mmol/L glucose. T2DM cells were also shown for comparison. (**A**) Up-regulated differentially expressed genes; (**B**) down-regulated differentially expressed genes.

Using a microarray dataset (GSE26168) from the Gene Expression Omnibus database, Zhu et al. identified 981 DEGs, of which 301 were upregulated and the rest downregulated^59^. These DEGs were highly enriched in cell differentiation, cell adhesion, intracellular signal transduction, and regulation of protein kinase activity, as well as cAMP signaling pathway, Rap1 signaling pathway, regulation of lipolysis in adipocytes, PI3K-Akt signaling pathway, and MAPK signaling pathway. Based on the PPI network of these DEGs, the top 6 genes contributing to T2DM initiation, progression, and intervention strategy were identified as PIK3R1, RAC1, GNG3, GNAI1, CDC42, and ITGB1. While none of these specific genes were in measurable levels in our cultures, GNG2, CDC20, CDCP1, and ITGB8 were upregulated, whereas PIK3IP1 and ITGB3 were downregulated in T2DM cells.

### Significant correlations between mRNA and protein expression levels

We conducted a comparative analysis (**Figures 5, 6**) to understand the relationship between mRNA and protein expression levels by comparing gene expression data obtained from RNA-seq with protein levels determined through mass spectrometry (MS). Proteins were isolated from the extracellular matrix that was synthesized by SMCs and deposited in the culture wells (ECM), as well as from the cytosol (C) and cytoskeleton (CS) of cells. We observed a moderate yet statistically significant correlation between mRNA abundance as estimated by RNA-seq, and protein abundance as determined from MS. This correlation was quantified using Spearman’s correlation coefficient (Rho), which ranged from 0.24 to 0.6 across various samples. These findings suggest that alterations at the transcriptional level are broadly reflected at the protein level, influencing biological pathways.

**Figure 5.**
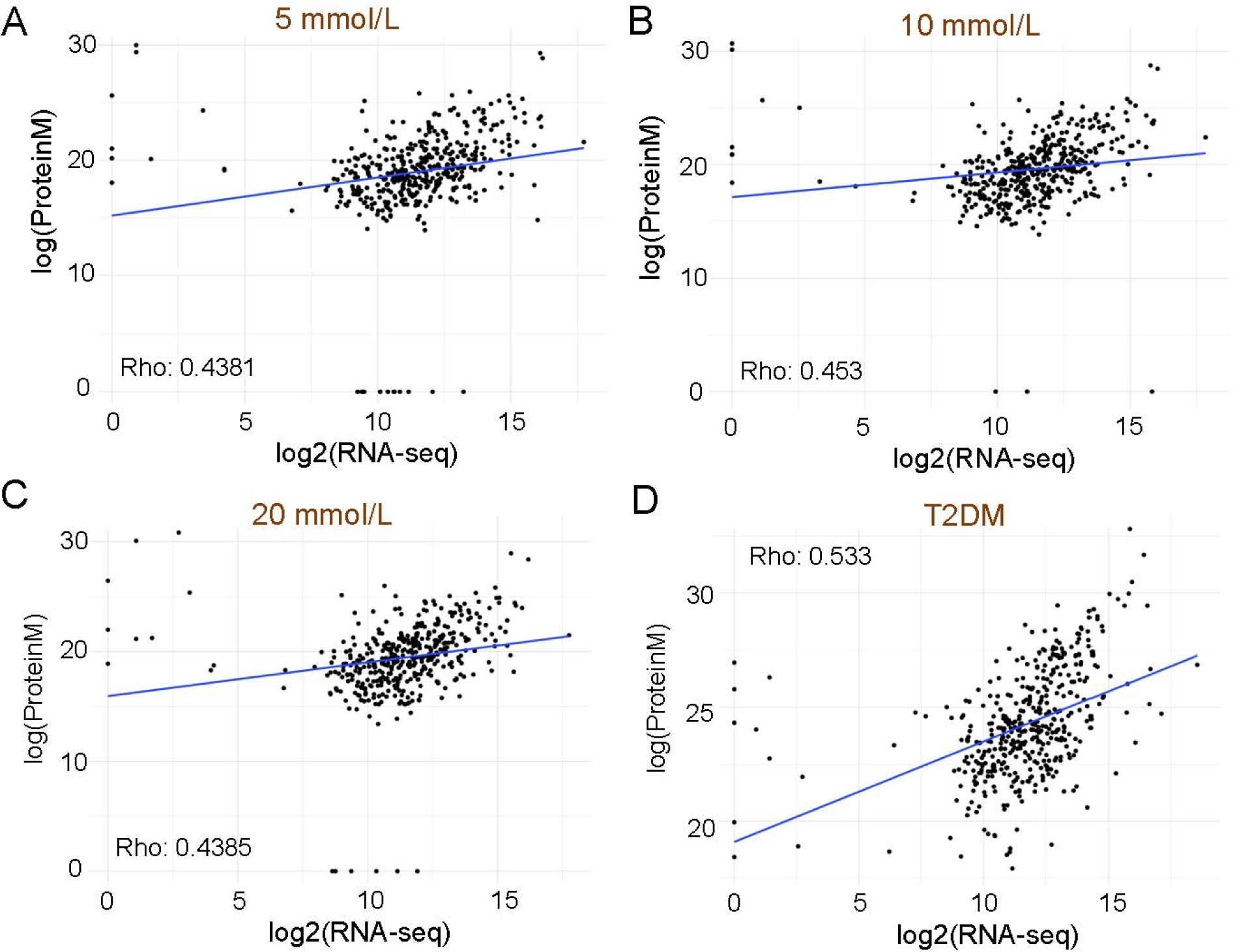
Correlation between ECM proteins and mRNA levels across various glucose conditions and T2DM patient samples for 448 genes. **A.** 5 mmol/L: Spearman correlation coefficient of 0.438 (*p* = 1.67×10^−21^). **B.** 10 mmol/L: Spearman correlation coefficient of 0.453 (*p* = 4.11×10^−23^). **C.** 20 mmol/L: Spearman correlation coefficient of 0.439 (*p* = 1.53×10^−21^). **D.** T2DM: Spearman correlation coefficient of 0.533 (*p* = 7.55×10^−33^).

**Figure 6.**
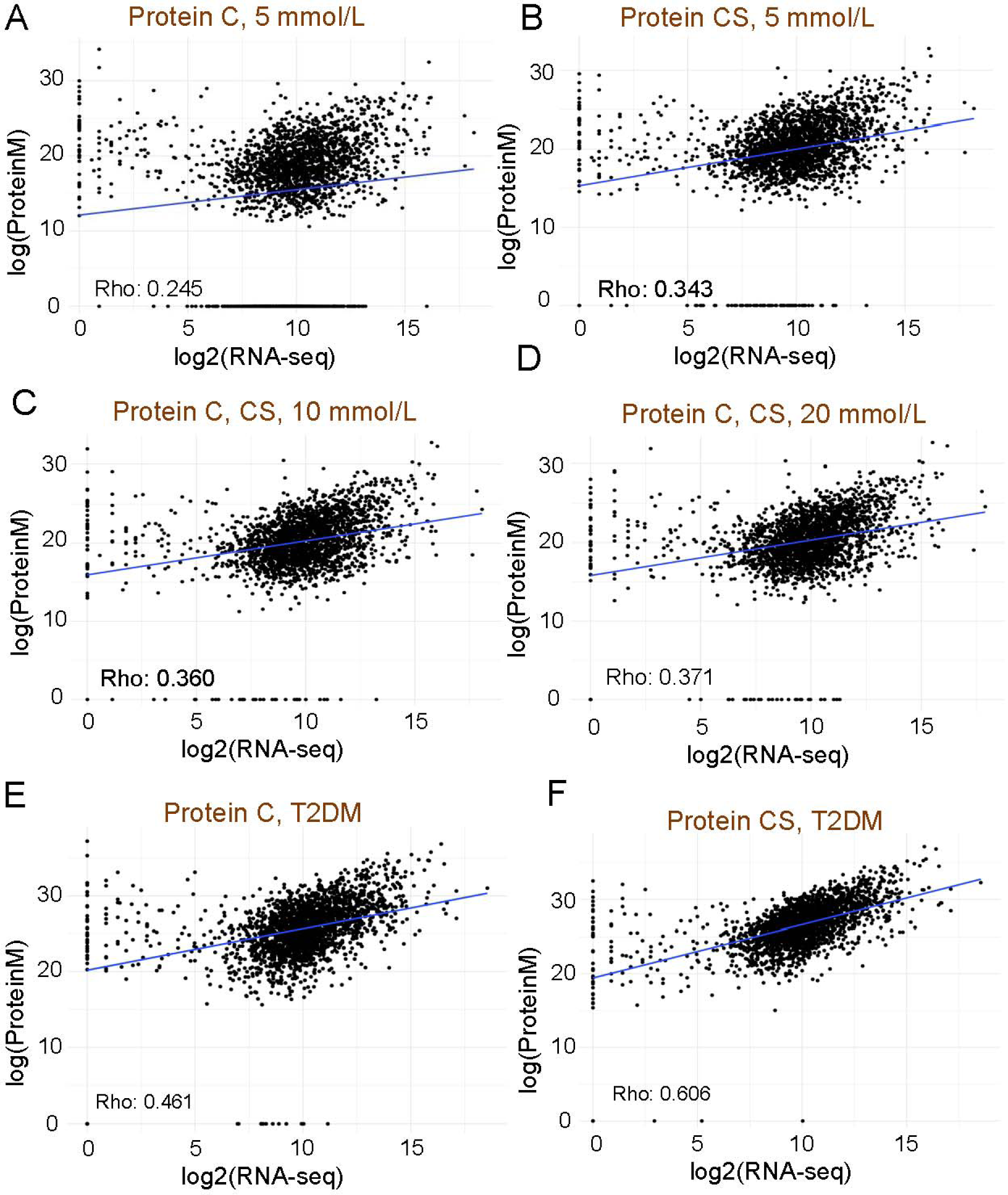
The correlation between mRNA expression levels and cytosol (C) / cytoskeletal (CS) protein levels for 2323 genes across different culture conditions. **A.** 5 mmol/L glucose: Spearman correlation coefficient of 0.245 (*p* = 2.67×10^−33^) for cytosol protein. **B.** 5 mmol/L condition: Spearman correlation coefficient of 0.343 (*p* = 3.71×10^−65^) for cytoskeleton protein. **C.** 10 mmol/L: Spearman correlation coefficient of 0.363 (*p* = 2.61×10^−72^) for both cytosol (C) and cytoskeleton (CS) protein. **D.** 20 mmol/L: Spearman correlation coefficient of 0.371 (*p* = 7.58×10^−77^) for both cytosol (C) and cytoskeleton (CS) protein. **E.** T2DM: Spearman correlation coefficient of 0.461 (*p* = 1.19×10^−122^) for cytosol protein. **F.** T2DM patient samples: Spearman correlation coefficient of 0.606 (*p* = 7.57×10^−234^) for cytoskeleton protein.

### Identifying metabolites co-expressed genes and enriched pathways

Gene expressions and metabolites constitute two distinct layers of features that can characterize dynamic responses to changes in glucose levels. We investigated which genes are co-expressed with metabolites. For each metabolite, we identified gene sets that exhibited similar dynamic patterns, using Spearman’s rank correlation coefficient (Rho) with a threshold of |Rho| > 0.9. Subsequently, we conducted a pathway enrichment analysis to determine which pathways are enriched in these co-expressed gene modules, which show similar patterns to the metabolites.

Notably, the L13a-mediated translational silencing pathway and the neutrophil degranulation pathway were repeatedly found to be enriched, linked to genes co-expressed with 9 and 7 different compounds respectively, as depicted in **Figures 7 and 8**. The L13a pathway plays a crucial role in cellular stress responses and could potentially reduce inflammation triggered by elevated glucose levels by regulating the translation of pro-inflammatory cytokines, thus acting as a defense mechanism against stress induced by hyperglycemia. Meanwhile, the neutrophil degranulation pathway, crucial for immune defense through the release of enzymes and reactive substances, might aggravate tissue damage under hyperglycemic conditions by promoting neutrophil activation and degranulation. These findings suggest that metabolites induced by hyperglycemia can indirectly indicate immune response signals, as inferred through pathway enrichment analysis using co-expressed genes with metabolites. A comprehensive list of enriched pathways associated with each metabolite’s co-expressed genes is presented in **Table S2**.

**Figure 7.**
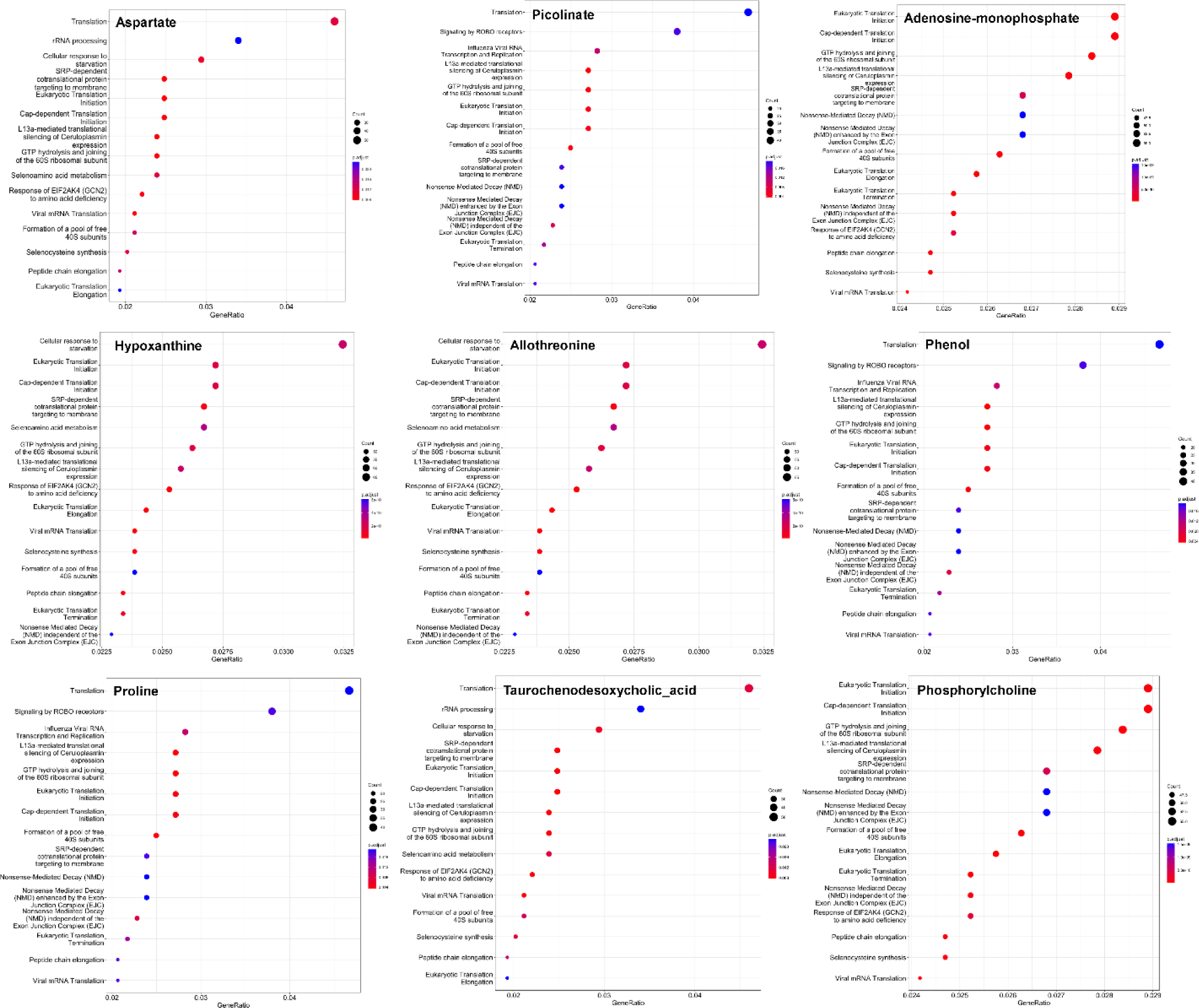
**Nine enriched pathways from co-expression analysis associated with compounds involving L13a-mediated translational silencing of ceruloplasmin expression**: aspartate, picolinate, adenosine-monophosphate, hypoxanthine, allothreonine, phenol, proline, taurochenodesoxycholic acid, and phosphorylcholine.

**Figure 8.**
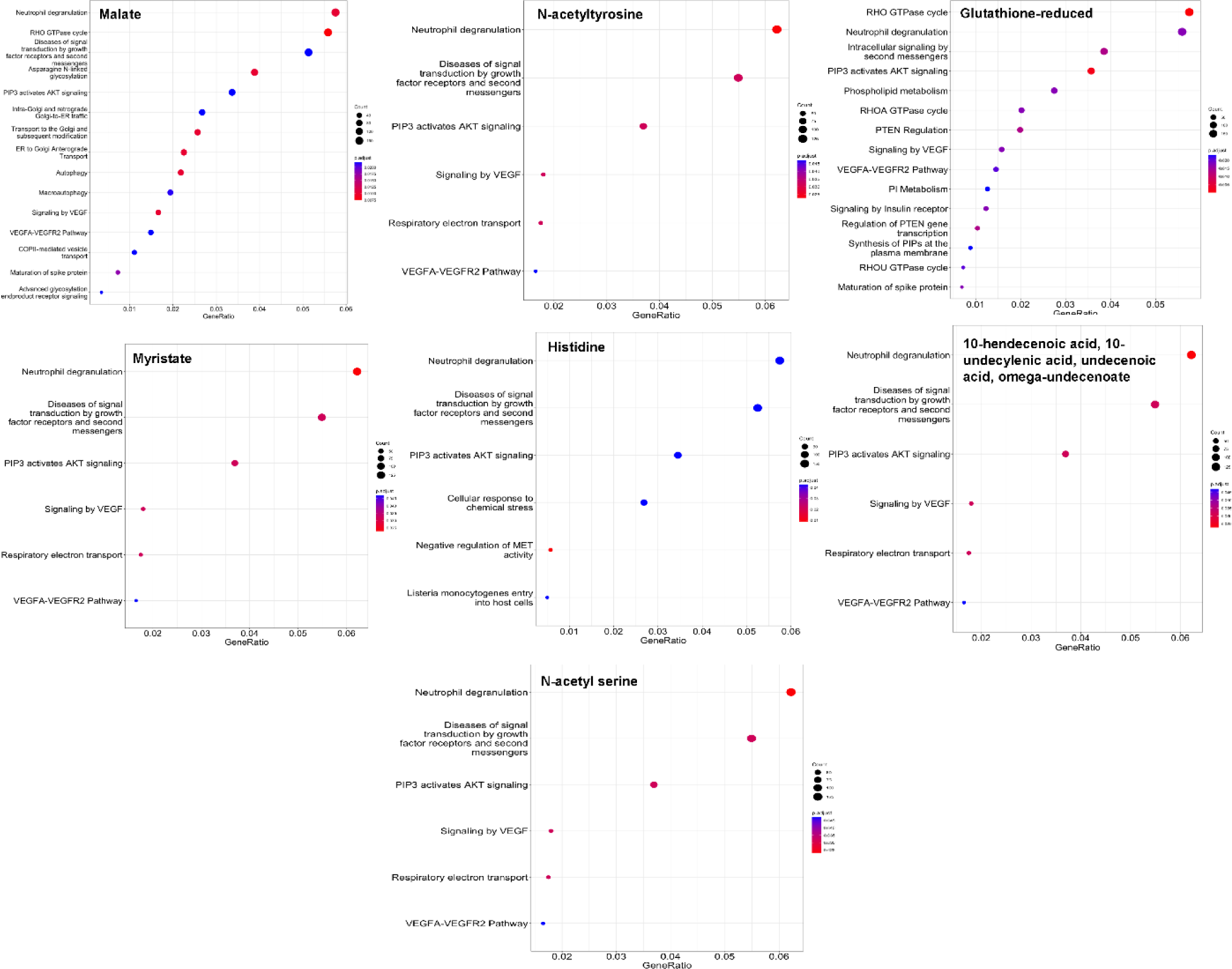
Seven enriched pathways from co-expression analysis associated with compounds involving Neutrophil degranulation: Malate, N-acetyltyrosine, glutathione reduced, Myristate, Histidine, undecenoic acid, and N-acetylserine.

Among the compounds involving L13a-mediated translational silencing of ceruloplasmin expression (Fig. 7), proline and picolinate are interesting. For instance, proline elevation is associated with lactic acidosis, and plasma proline levels strongly correlate with hemoglobin A1c and insulin-related variables (e.g., C-peptide, insulin, HOMA-IR)^60^, although dietary proline intake was not significantly associated with T2DM in and adult cohort study^61^. Picolinic acid, on the other hand, is an efficient chelator, and chromium picolinate has been promoted as a dietary supplement for T2DM patients owing to its role in blood glucose regulation, carbohydrate and lipid metabolism, and body composition^62^.

In a similar vein, prior studies have shown that high glucose and inflammatory conditions coax primary aortic SMCs to express higher levels of G6PDH and NADPH oxidase, elevated pentose phosphate pathway activity, and upper glycolysis pathway^63, 64^. Other pathways that are enhanced in SMCs under hyperglycemic conditions include protein kinase C signaling, RAGE (receptor for advanced glycation end products)/ERK/NF-κB signaling pathway, and ERK/Akt signaling pathway, which contribute to increased formation and accumulation of AGEs, progression of atherosclerosis and vascular calcification, induction of iNOS activity, autophagy, and VSMC proliferation, among others^65^.

### Limitations of this study

While our study extends human SMC exposure to hyperglycemia *in vitro* from acute (typically < 72 h) to chronic (3-week culture) culture, diabetes develops over a long period *in vivo*, sometimes over decades, which would be hard to replicate *in vitro*. T2DM is a complicated process involving interactions between multiple cell types, and hyperglycemia is perhaps just one of the contributors to this condition, which warrants further *in vitro* investigations on other contributors. For these findings to be relevant to human T2DM, selected studies in animal models are required to elucidate the mechanisms involved and identify possible therapeutic applications. Furthermore, our *in vitro* results should be validated in other healthy and T2DM patient derived SMCs *in vitro*, and *in vivo* in future studies.

### Conclusions

This study investigates the effects of hyperglycemic condition on human aortic SMCs cultured under varying glucose concentrations (5 mmol/L, 10 mmol/L, 20 mmol/L), and compared to SMCs isolated from patients with type 2 diabetes (T2DM). We noted that chronic hyperglycemia, in a glucose concentration dependent manner, induced significant changes within human SMCs *in vitro* in their (i) phenotype and biophysical characteristics, (ii) biomechanical properties (Young’s modulus, membrane tension, tether and adhesion forces), (iii) cytokines/ chemokines/ growth factors/ and MMPs-TIMPs release, and (iv) differential expressions in gene-sets and pathways representing inflammation, matrix turnover, and metabolic pathways. We believe that the biomechanical results (*F_T_*, *F_ad_*, *T_M_* and *R_T_*) presented here for human T2DM cells and those of human SMCs under hyperglycemic conditions haven’t been reported earlier in literature. Finally, our integrated analysis of omics datasets revealed specific biomarkers and enriched pathways, which we report for the first time on T2DM cells and on human SMCs receiving higher glucose levels, furthering our knowledge on chronic hyperglycemia-induced changes and their complex relationships.

## Declarations

### Ethics approval and consent to participate

Not applicable

### Consent for publication

Not applicable

### Availability of data and material

The datasets generated and/or analyzed during the current study are available from the corresponding authors on reasonable request.

### Competing interests

The authors declare that they have no competing interests.

### Funding

NHLBI/NIH Progenitor Cell Translational Consortium (PCTC) (MIRC-002500: to PJ) and Defense Advanced Research Projects Agency (DARPA) (AWD00001593: to PJ); Partial support from National Science Foundation (NSF) grants 1927602 and 1337859 to CK; and the Cellular and Molecular Medicine Fellowship from Cleveland State University to SB.

### Author Contributions

CK: conceptualization, formal analysis, funding acquisition, project administration, supervision, writing original draft, review and editing. PJ: funding acquisition, computational analysis, methodology, supervision, writing original draft, review and editing. SB: experimental design, formal analysis, methodology, writing original draft. AB: computational analysis, methodology. YS: methodology, metabolomics, supervision, manuscript editing and review. IR: metabolomics, experimental setup and data analysis. EGE: atomic force microscopy, data acquisition and analysis.

## Acknowledgements

We acknowledge access to the Mass Spectrometry facility in the Department of Chemistry at Cleveland State University. We would like to express our gratitude to the Genomics Core at Cleveland Clinic for their invaluable assistance with library preparation and sequencing.

## Supplementary Information

The online version contains supplementary material.

## Abbreviations

AFM: Atomic force microscopy
T2DM: Type 2 diabetes mellitus
CVD: cardiovascular diseases
VEGF: vascular endothelial growth factor
NO: nitric oxide
VSMCs: Vascular smooth muscle cells
ECs: Endothelial cells
GAGs: glycosaminoglycans
IGF: Insulin-like growth factor
ECM: Extracellular matrix
ROS: Reactive oxygen species
NF-kappaB: nuclear factor kappaB
RAGE: Receptor for advanced glycation end-products
FAK: Focal adhesion kinase
MMP-2: Matrix metalloproteinase-2
ICAM-1: Intercellular cell adhesion molecule-1
VCAM-1: Vascular cell adhesion molecule-1
HAOSMC: Human aortic smooth muscle cells
T2DM-HAOSMCs: Type II diabetic patients derived aortic smooth muscle cells

## Notes

### Competing Interest Statement

The authors have declared no competing interest.

